# Spatio-temporal structure of single neuron subthalamic activity in Tourette Syndrome explored during DBS procedures

**DOI:** 10.1101/532200

**Authors:** Matteo Vissani, Roberto Cordella, Silvestro Micera, Luigi M. Romito, Alberto Mazzoni

**Author notes:** these authors share equal senior contribution (Luigi M. Romito, Alberto Mazzoni).

## Abstract

Basal ganglia dysfunctions have been suggested to play a causal role in the pathophysiology of most motor and non-motor symptoms of movement disorders as Tourette Syndrome (TS) or Parkinson’s Disease (PD). Intra/post-operative recordings from the subthalamic nucleus (STN) during Deep Brain Stimulation (DBS) procedures in PD patients have highlighted specific pathological patterns of neural activity. Spatial and temporal patterns of STN neural activity in TS are still unknown due to the lack of direct microrecordings in humans. Here, we describe for the first time specific neural activities of sensorimotor STN in TS patients, as recorded during intraoperative microrecordings. We analyzed 125 single units at 0.5 mm-spaced depths from the STN of anesthetized TS patients and we observed a large fraction of units (39/125, 31.2%) intensely bursting in the delta band (<4 Hz). In anesthetized PD patients we found similar average firing rate and spectral density of STN units, but differently to TS patients, only 4/54 (7.4%) of the units displayed bursting. Remarkably, bursting units in TS STN were not homogeneously distributed over the dorso-ventral trajectory of the recording: the highest density of bursting units was reliably found at the depth for which the clinical effect was maximal. Our results provide an unprecedented characterization of STN functional architecture and single units dynamics in TS patients, paving the way to an understanding of the role of STN subterritories in TS.

**Key Points:**

- Single neuron activity in Subthalamic Nucleus (STN) of patients with Tourette Syndrome (TS) was analyzed for the first time in literature.
- Firing rate and spectral content of single STN neurons in TS patients were found to be similar to those of anesthetized PD patients, while the analysis of arrhythmic bursting activity revealed that in TS patients the STN is characterized by a larger fraction of bursting neurons and more intense bursts
- Bursting activity in TS was widespread across the whole STN, but with a higher density at the optimal lead location depth for DBS

## Introduction

Tourette Syndrome (TS) is a childhood-onset neuropsychiatric disorder characterized by motor and vocal tics (American Psychiatric Association, 2013;Robertson *et al.*, 2017). In the last two decades, a number of studies associated TS tics to a dysfunctional interaction between the motor cortex and the basal ganglia (BG) stemming from an aberrant behavior of the latter (Albin, 2006;Bronfeld & Bar-Gad, 2013;Yael *et al.*, 2015). The BG is a complex system sending feedbacks to the cortex through the thalamus composed by six different elements: the STN and the Striatum receive cortical inputs, the substantia nigra compacta (SNc) and the external globus pallidus (GPe) are internal processing units, and the internal globus pallidus (GPi) and the substantia nigra pars reticulata (SNr) are output stations projecting to the thalamus, which in turn modulates cortical activity (Albin & Mink, 2006;Lanciego *et al.*, 2012;Mink, 1996;Mink, 2006;Utter & Basso, 2008). The functional interplay among these elements is given by the combination of three loops. In the direct pathway (DP) the Striatum is excited by the cortical activity and inhibits the GPi/SNr complex. This reduces inhibition to the thalamus, increasing then cortical activity. The DP is then a positive feedback loop reinforcing movements triggered by the motor cortex. In the hyperdirect pathway (HP) the STN is excited by the cortex and excites the GPi/SNr complex leading to a reduction of thalamic excitation to the cortex. This makes the HP a negative feedback loop. In the indirect pathway (IP) the striatum inhibits the GPe, which reduces the inhibition toward the GPi/SNr complex and directly increases GPi activity through STN excitation. The net result is an inhibition of thalamic and hence cortical activity and an overall negative feedback loop.

The widely accepted hypothesis about the origin of TS disorders is that they might originate from an overactivation of the striatum, generating an abnormal reinforcement of the DP and hence an excessive positive feedback of motor cortical activity leading to involuntary movements (Bronfeld & Bar-Gad, 2013;Bronfeld *et al.*, 2013;Mink, 2001a). Support for this hypothesis derives from recordings in nonhuman (Bronfeld *et al.*, 2011) and human (Ganos *et al.*, 2015) primates showing a relationship between the striatal region somatotopically associated with the tic-expressing body part and the probability of its occurrence. Striatum overactivation can have different causes, as reduction in the number of striatum interneurons (Kalanithi *et al.*, 2005;Kataoka *et al.*, 2010) or functional disinhibition due to dopamine circuits’ dysfunctions (Crossman *et al.*, 1988;Singer & Minzer, 2003;Worbe *et al.*, 2013). Limitations of this model are that it does not account for any relevant involvement of the STN (Yael *et al.*, 2015), and that it largely relies on animal experiments.

TS treatment is often hard to be efficaciously personalized (Efron & Dale, 2018;Hartmann & Worbe, 2018) and should aim to ameliorate tics and behavioral disorders. In pharmacologically refractory TS patients, however, basal ganglia or thalamic deep brain stimulation (DBS) has emerged as a promising treatment (Baldermann *et al.*, 2016;Pena *et al.*, 2018). In the past twenty years DBS treatment of TS demonstrated to be useful and efficient therapy for many pharmacologically-resistant and behavioral-refractory TS patients (Baldermann *et al.*, 2016;Servello *et al.*, 2010;Shahed *et al.*, 2007;Vandewalle *et al.*, 1999). DBS-driven modulation of the thalamic and GPi oscillatory activity has been correlated to clinical outcome (Maling *et al.*, 2012), but the effect of DBS stimulation of STN in TS patients has never been properly investigated (Baldermann *et al.*, 2016).

In addition to a direct therapeutic role, the advent of DBS provided a unique and powerful platform for the investigation of the neuropathophysiological mechanisms of TS. Indeed, DBS studies contributed to validate the DP-enhancement model of the origin of TS (Hashemiyoon *et al.*, 2017;Israelashvili *et al.*, 2015;Priori *et al.*, 2013), revealing in the excessive theta local field potential activity and in the theta synchronization between GPi and thalamus a hypothetical hyperkinetic neuropathological marker (Neumann *et al.*, 2018).

In this study we contribute to shed light on the dynamics of STN in TS through the analysis of intraoperative microrecordings performed for the very first time in the STN and subthalamic region. These neurophysiological observations were made while conducting a phase II study on efficacy and safety of STN DBS in treating motor and phonic tics in medically refractory pure (motor) – TS patients (ClinicalTrials.gov ID: NCT02619084, research funded by the Italian National Institute of Health (NIH), GR-2009-1594645, PI L. Romito).

Due to the patient’s condition these recordings were performed during surgical procedure and microrecording on TS patients under general anesthesia. To check for the effects of the recording procedures on the observed STN activities, the results were compared with recordings performed on PD patients of ours, who underwent STN implant under general anesthesia due to excessive rest tremor. Results of these compared analysis highlighted TS-specific neural dynamics in STN, in particular neurons generating low frequency arrhythmic bursts, with a non-homogeneous depth distribution.

## Methods

### Ethical Approval

The study was approved by the Carlo Besta Institutional Ethics Committee (reference number 7/2014) and was carried out according to the Declaration of Helsinki. Informed written consent was obtained either from the patients or from their legal representative.

### Subjects

Inclusion criteria for the TS patients were based on current diagnostic criteria (American Psychiatric Association, 2013): age of at least of 18 years with a stable degree of severity and no spontaneous improvement of tics, a Yale Tic Global Severity Scale (YTGSS) > 35/50 for at least 12 months, with tic severity documented by a standardized videotape assessment, inadequate response to standard drug treatment (medication or botulinum toxin).

The following exclusion criteria were implemented: diagnosis of secondary tic disorder, cognitive or severe psychiatric abnormalities.

Intraoperative microrecording data were compared to those obtained from two parkinsonian patients who underwent STN implant procedure under general anesthesia, and who were enrolled for DBS according recommendations of the CAPSIT-PD panel (Defer *et al.*, 1999).

### Anesthesia and Surgical Procedure

Surgery was performed, for both TS and PD patients, with the BIS-Guided general anesthesia through Target Controlled Infusion (TCI) of Propofol and Remifentanil. Bispectral Index (BIS), a depth of anesthesia (DOA) monitor, might provide an electroencephalography (EEG) correlation with human behavior in the course of general anesthesia surgeries. BIS value specifies the likelihood of consciousness, thus aiding the tailoring of anesthetic dosages to the individual patient. Anesthetics delivery was targeted to keep BIS values between 40 and 60 (Cordella *et al.*, 2018).

Surgery was always performed in stereotactic conditions with the Leksell (Elekta Inc., Atlanta GA, USA) frame. Computerized tomography (CT) was used to recognize the anterior and posterior commissures in the stereotactic space, after positioning of the Leksell-modified headframe. A probabilistic stereotactic digitalized atlas registered to the anterior commissure (AC)–posterior commissure (PC) line was utilized for the preliminary determination of the target coordinates. The stereotactic CT images were then merged with the preoperative magnetic resonance images (MR; T1 and fast spin echo inversion recovery sequences with double dose of contrast agent) and a second calculation of the target coordinates was performed with the neuronavigation system (Stealth Station Treon Sofamor Danek, Medtronic Inc. Minneapolis, MN, USA). Finally, the two set of data were matched, and the definitive coordinates, along with the planned trajectory, were adapted to the individual patient anatomy. A rigid cannula was then inserted through a small hand-drilled burr hole (5-mm diameter) and through the opened dura mater; the cannula was then indwelled till 15 mm above the estimated target. A high impedance microelectrode (250-lm tip, and impedance 1–1.5 MX; FHC Inc., Bowdoinham ME, USA) was then introduced within the cannula and advanced progressively to the target. If microrecording fulfilled the localizing criteria, the definitive electrode (Leads Model 3389, Medtronic Inc. Minneapolis, MN, USA) was placed at the target through the same rigid cannula after the microelectrode withdrawal. Macrostimulation was then performed in bipolar configuration between the two proximal contacts and between the two distal contacts. At this stage of the procedure, clinical testing of the awake patient and electromyographic (EMG) recording in the anesthetized patient allowed to search for favorable clinical response and/or for the appearance of side effects. If microrecordings and macrostimulation through the definitive lead suggested its accurate placement at the right target, the lead was secured to the burr hole with fibrin glue and a titanium plate. Otherwise if microrecordings and/or clinical and EMG monitoring suggest a suboptimal electrode placement, a new trajectory was planned through the same burr hole. Each definitive implanted lead was finally connected to a Medtronic Activa® SC 37603 Neurostimulator through a Medtronic Model 37086 Extension.

The location of the therapeutic contacts and the volume of tissue activated by the electrical field around them was ascertained by Medtronic StealthStation TREON plus Navigation System. Three-dimensional anatomy of basal ganglia was adapted to the brain geometry of each patient by overlaying the pre-operative and postoperative MRI or computed tomography scan onto the software atlas. The volume of activated tissue was estimated by using the contact(s) and the electrical parameters set when the patient had reached a stable clinical condition. Active contacts location was also reconstructed by using LEAD-DBS v2.1.8 (http://www.lead-dbs.org/) (Horn & Kuhn, 2015;Horn *et al.*, 2019).

Electrodes were considered as correctly positioned only if the electrode contacts, the electric field volumes around the stimulated contacts, or both, included the STN. These procedures were done by a neurosurgeon blinded to the results of neurostimulation in each patient.

### Electrophysiological recordings

Continuous physiological recordings began at 10 mm above the target and were performed by means of a Medtronic LeadpointTM system (Medtronic Inc., Minneapolis, MN, USA). An exploratory trajectory was made by extruding the microelectrode (250 lm tip, and impedance 1–1.5 MX; FHC Inc., Bowdoinham ME, USA). Microrecording tracks are performed with 0.5 mm steps, and 1\2 mm beyond the target along the single planned trajectory. In order to investigate the neuronal response properties in well isolated neurons, stimuli such as light touch, pressure to the skin, pin prick and passive movements were systematically delivered to the whole body and contralateral, to the recording site, arms and legs (Cordella *et al.*, 2010).

### Spikes detection and sorting

The signals were exported at 24 kHz and transformed into .mat files for off-line analysis with custom MATLAB (Mathworks) routines. For the single unit analysis, every spiking period that was corrupted by noise or by spike amplitude instability that could affect the analysis was visually identified and discarded, i.e. a track was considered reliable if the noise period was below the 50% of the total length of the signal.

Spike detection and sorting were performed using Wave_Clus (Quiroga *et al.*, 2004), the most popular free-downloadable MATLAB ToolBox at this purpose. Briefly, the steps typically involved in spike detection and sorting are: (i) band-pass filtering the data, for example between 300 and 3000 Hz, using a zero-phase filter to avoid distortion in the spike waveforms (Quian *et al.*, 2009); (ii) computation of a double (positive and negative) detection threshold based on a robust estimate of the noise statistics; (iii) feature extraction of the spike shapes, using the wavelet transform; and (iv) clustering of the waveforms to identify the firing of the different units by means a modified k-means algorithm.

Every isolated unit was visually inspected and only the well separated ones were selected for further analysis. We used the following criteria (Giorni *et al.*, 2017): more than 90% of the total area of the amplitude histogram had to be above the detection threshold; the mean waveform had to have a typical action potential shape, i.e. biphasic shape either with a positive or negative most prominent peak; the percentage of spikes occurring within 3 ms of each other had to be less than 1% and the number of spikes detected had to be more than 20.

### Firing rate estimation

Firing rates of the neurons were estimated using the Gaussian Kernel bandwidth optimization method, a classical procedure for estimating the firing rate (FR). It has been shown (Shimazaki & Shinomoto, 2010) that the classical kernel methods accomplish well in the goodness-of-fit to the underlying rate and they outperformed competing methods in representing abrupt changes, i.e. non-stationarities, in the spike rate, which we often encounter in neuroscience. The kernel function *k*_*w*_ is a Gaussian function with mean value t_i_ the i-th spike timestamp and deviation standard w the bandwidth that is optimized minimizing a cost-function C_w_, according the following equations:

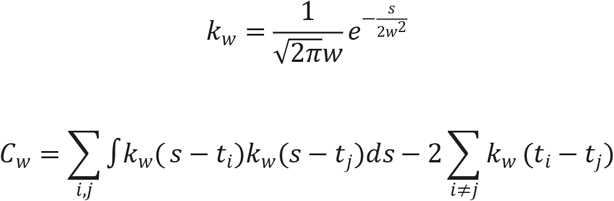

### Discharge pattern analysis and burst detection

The firing regularity was quantified by fitting the ISI distribution with a Gamma distribution (Mochizuki *et al.*, 2016):

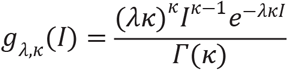

where I is the duration of a given ISI, κ and λ are the firing rate and scale factor, and *Γ*(*κ*) is the gamma function. The shape factor represents the firing regularity itself and it is estimated by the maximizing the likelihood, which is achieved by the following equation:

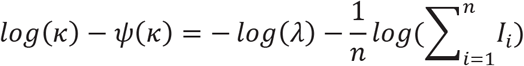

where *Ψ*(*κ*) and n are the digamma function and the number of ISIs respectively. Firing regularity log κ is expected to take a value of 0 (i.e., κ = 1) for a Poisson-like irregular spike train and becomes positive or negative if firings are more regular (κ > 1) or bursting (κ < 1), respectively. We used *log*(*κ*) as a criterion to determine three distinct patterns: for *log*(*κ*) > 0.3 the neuron was automatically classified as tonic. Instead, for 0 > *log*(*κ*) > −0.3 we used an additional criterion based on the firing rate profile: if > 70% of the samples was outside the range *λ* ± 0.5*λ*, i.e. presence of periods with very high and low firing rate, the neuron was classified as burst-like, otherwise as irregular.

The rank surprise method (Gourevitch & Eggermont, 2007) was used to identify a burst (intuitively, an occurrence of many spikes in a small time interval), in the spike train of the burst-like labelled cells. This method has a structure like the Poisson method (Legéndry & Salcman, 1985) and produces comparable results. Instead of assuming a Poisson ISI distribution, the Rank surprise assigns a uniform probability to each ISI, based on its rank, and proposes a different statistic, the rank statistic (RS), to evaluate a burst. We need to specify two parameters: the largest ISI allowed in a burst (limit), e.g. the third quartile as suggested in literature, and the minimum RS (threshold) that a burst must have to be considered valid. First, we search the spike train for all the ISIs that are smaller than limit and mark them. Then, we compute the RS value for every possible continuous sequence (time period) of marked ISIs and find the one with the largest RS. If this RS value is larger than the threshold, we denote a burst and remove the corresponding ISIs from the marked ones. Finally, we repeated the process for the remaining marked ISIs. Furthermore, we computed five different parameters of bursts: (1) the mean intra-burst frequency (IBF) as the number of the spikes within burst divided by the burst length (2) the mean inter-burst interval (IBI) as the time interval between two consecutive bursts, (3) the mean spikes number within bursts (SWB), (4) the proportion of spikes in bursts (SIB) during 10 s of the recording and (5) the coefficient of variation (CV) as the standard deviation of the IBI divided by its mean.

### Oscillatory strength in neural activity

A method that relies on the estimation of the frequency spectrum from the normalized autocorrelation histogram (ACH) was used to investigate the neural oscillations at the single-neuron level (Muresan *et al.*, 2008). This method is not computationally demanding, minimizes the false detection of oscillations and can be used even with single units. Briefly, the steps typically involved in this procedure are: (i) a fast Gaussian Kernel to smooth the ACH and to remove high frequency noise; (ii) a slow Gaussian Kernel to detect the boundaries of the ACH central peak; (iii) efficient removal of the central peak from the smoothed ACH; (iv) the peak less smoothed ACH is subjected to FFT; (v) peak detection within the band of interest. Eventually, a parameter which quantifies the oscillation’s strength called oscillation score *ϑ*_*s*_ is computed as the ratio between the highest frequency magnitude within the band of interest and the baseline, e.g. the average, magnitude of the spectrum. To obtain a measure of the confidence on the oscillation score in a single-trial and few samples context as our case, we defined “reliable” oscillation scores that were greater a threshold determined by the mean *μ* and the standard deviation σ of the total spectrum, as follows:

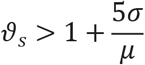

Prevalence of units with significant oscillatory was then assessed for each of the following frequency bands: delta [1-4] Hz, theta [4-8] Hz, alpha [8-12] Hz, beta [12-30] Hz and gamma [30-100] Hz.

### Mutual information analysis

We computed the mutual information about the target depth carried by three neural properties: i) the presence of bursting neurons, ii) the presence of oscillatory behaviors and iii) the firing rate, using the Information Breakdown Toolbox in Matlab (Magri *et al.*, 2009).

Mutual information between a set of responses R and a set of neural signals S is defined as (Shannon, 1948):

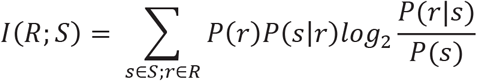

where P(r) and P(s) are the absolute probability across all trials (neurons in our case) of observing a given response r from the set R or given neural signal s from set S, and P(s|r) is the conditional probability of observing the neural signal s in trials (neurons in our case) in which the response is r. Here, we considered as set of responses the relative target-depth of the neurons: before (more dorsal), on and after (more ventral) the target position and as neural signals the firing rate (low firing rate and high firing rate according to the median value), the oscillation presence (1: oscillatory; 0: not-oscillatory) and the presence of bursting neurons (1: bursting neuron; 0: not-bursting neuron). We tackled the information bias due to the small data set (Panzeri *et al*., 2007) with the following precautions: i) we limited the number of bins of the neural signals to two to ensure a conservative but stable measure of information (Ince *et al.*, 2003); ii) we applied the Panzeri-Treves bias correction (Treves & Panzeri, 1995) and iii) we compared the resulting values of information with those obtained with 500 bootstrap repetitions using P < 0.05 as an information significance threshold.

### Statistical Tests

JMP 14.0.1 software (SAS Institute Inc. Cary, US) and MATLAB R2017b (Mathworks, Natick, MA, USA) custom-written scripts were used for statistical analysis. Fisher-Exact test was used to examine the significance of the association in contingency tables categorical data. A Shapiro-Wilk test was used to determine if the datasets were normally distributed. None of our datasets passed the test substantial inasmuch as a prominent right skewness was present in almost all distributions. Therefore, non-parametric statistics (Mann-Whitney U) tests were used for comparisons of neural activities in patients with TS or PD and, unless otherwise noted, all group values of continuous variables were reported as median ± half of the interquartile range (IQR). Linear regression and binning analysis were used to describe the relationship between neural features and depth. Pearson’s correlation coefficient was used to examine the linearity of the relationship and R squared to quantify the variance explained by the linear model. Results were considered significant at P < 0.05.

## Results

### Patients and DBS therapy

This work aims at offering a detailed characterization of the spatio-temporal structure of STN firing dynamics in subjects affected by TS. Four pure-TS (Eapen & Robertson, 2015) patients (2M/2F, age at implant 34.0 years, range 26-46) underwent bilateral STN implant (Medtronic 3389 lead, Activa SC pulse generator) at the Carlo Besta Neurological Institute (Table 1). Two PD patients were otherwise enrolled for data comparison.

**Table 1.** Baseline characteristics of 4 TS patients. Mean ± SD given in the last row where appropriate.

| **N patient** | **Gender** | **Age at implant (Y)** | **Disease duration (Y)** | **YTGSS tic total score** | **YTGSS impairment score** |
| --- | --- | --- | --- | --- | --- |
| **ID1** | F | 29 | 23 | 42 | 50 |
| **ID2** | M | 26 | 19 | 38 | 50 |
| **ID3** | F | 35 | 22 | 44 | 50 |
| **ID4** | M | 46 | 33 | 42 | 40 |
| **Tot** | 2F/2M | 34.0±8.8 | 24.3±6.1 | 41.5±2.5 | 47.5±5.0 |

Neural recordings were performed with microelectrodes during the exploratory recordings to determine the optimal target location for DBS (Figure 1A-B, Table 2, Methods for details). In all patients, an important subthalamotomic-like improvement of motor and phonic tics was observed soon after the implant and lasted on average 4.5±2.4 days. No dyskinesia was detected during this period. In the days and weeks after the clearing of this effect, we started the programming sessions. Initial DBS parameters were set as monopolar, 0-3 Volts (0.1 V steps), 125 Hz, 60 μs stimulation, starting with the ventral-most contact and then preceding towards the dorsal-most contact. In all patients, a sustained acute tic improvement was observed following stimulation of STN-located contacts, even at low stimulation current (average 0.6 ± 0.4 mA) (Figure 1C, Wilcoxon signed rank test, P < .001). This result proves that i) STN is an efficient target for reducing TS motor symptoms, ii) the location selected within STN was highly effective.

**Table 2.**
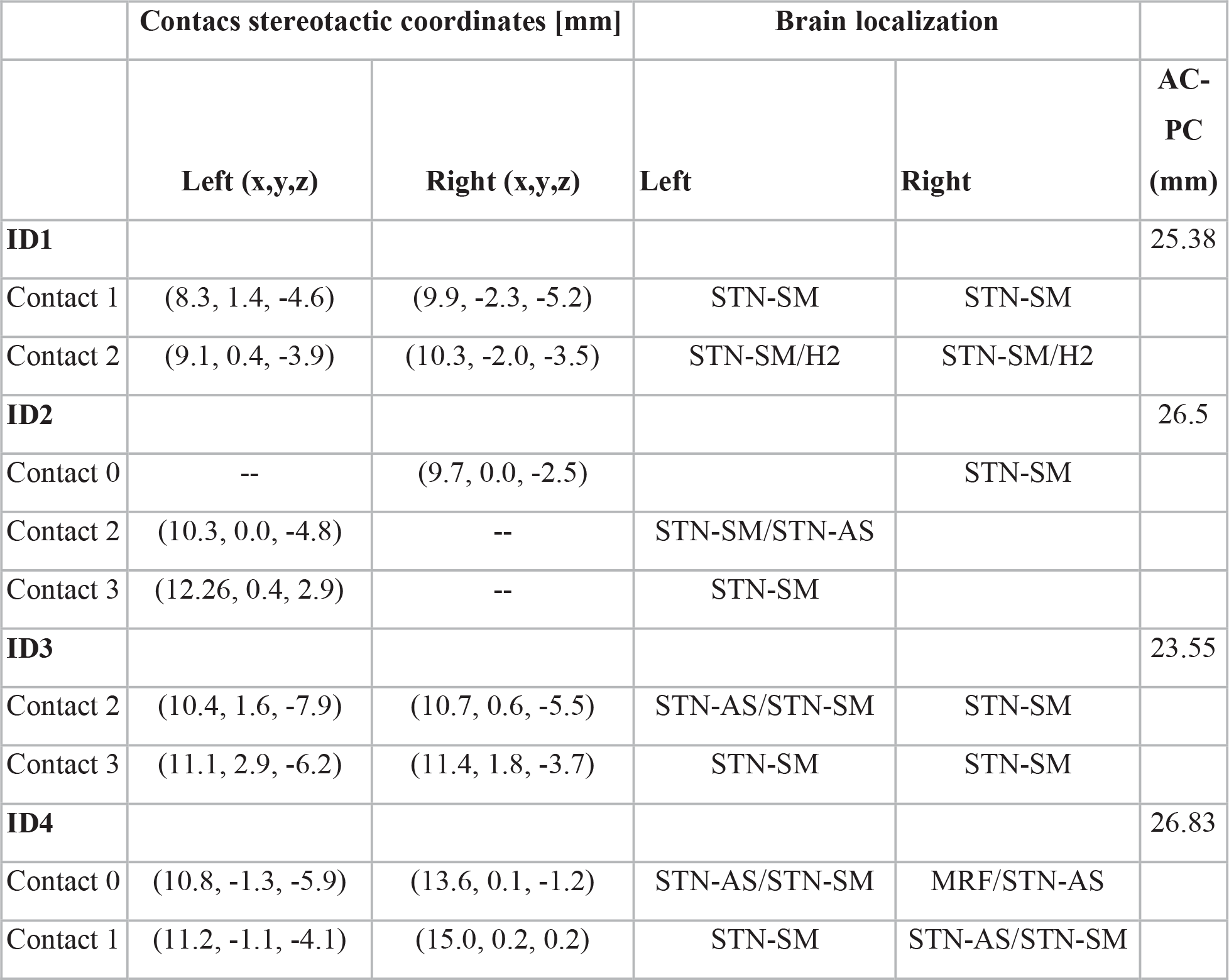
Stereotactic coordinates of active contacts from the post-operative CT-MRI fusion (TS patients). Brain localization is classified as STN Sensory-Motor (SM), Associative (AS), Forel’s Field H2 (H2) or Mesencephalic Reticular Formation (MRF).

**Figure 1.**
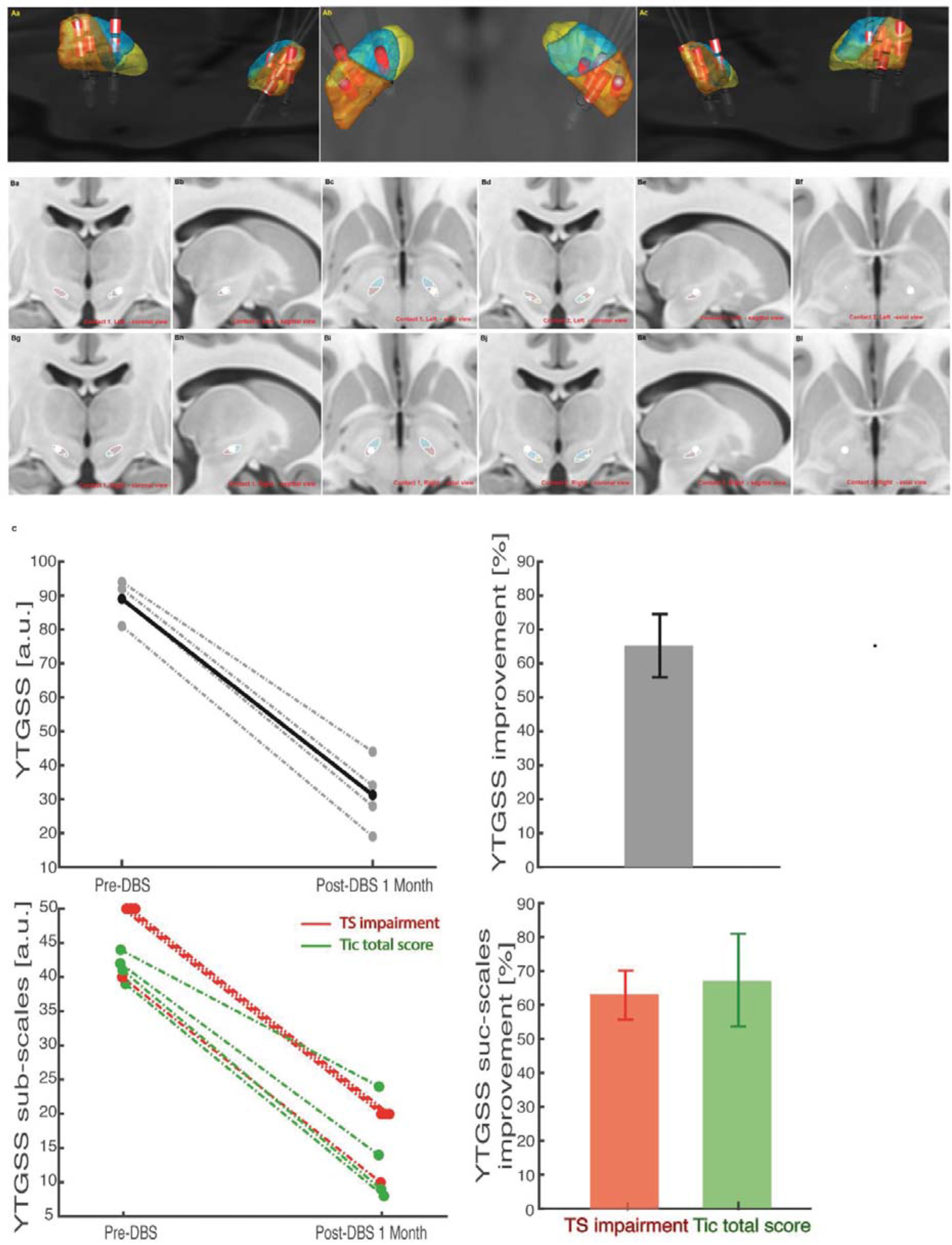
Surgical targeting, electrode placement (reconstructed using Lead-DBS v2.1.8) and resulting clinical scales outcomes before and after DBS. *A*, group analysis showing the 3D location of leads and stimulating contacts (in red) in all four TS patients. Color codes: dark orange, sensorimotor STN; light cyan, associative STN; light yellow, limbic. *B*, Reconstruction of electrode positioning in patient 1 according to functional STN subdivision by Accolla (Accolla *et al.*, 2014). *C*, One-month post-operative outcome (tic scores measured by YTGSS).

Due to the strong and unpredictable movements of TS patients this operation was performed in general anesthesia. Exploratory recordings from PD patients, which were also performed in general anesthesia due to patient’s involuntary movements, were also analyzed to identify TS-specific features. As animal model studies suggest that bursting activity might be associated to TS tics, we first analyzed the presence of spiking patterns in the STN of TS. These patterns were then compared with those observed in PD patients.

### Similarities in firing activity in TS and PD STN neurons

We identified in all recordings the activity of individual neurons (see Methods for details). The size of the set of neurons identified across the four TS patients was 32 (± 9), in the range 19-45, for a total of n=125 neurons, while 13 and 41 neurons were identified in the two PD patients’ recordings. We then computed the average firing rate and the ISI distribution for the 125 STN neurons recorded from TS patients and the 54 STN neurons from PD patients. Comparison of the mean discharge rate showed that there was no significant difference in the mean activity between the patient groups (TS: 11.72 ± 6.1 spikes/sec, PD: 12.23 ± 7.3 spikes/sec, Mann-Whitney test, P = 0.73).

We performed also a spectral analysis of the firing rate to characterize the presence of fluctuations over different timescales in the activity of TS STN neurons. Neurons displaying significant peaks in the power spectrum of their firing rate autocorrelation (see Methods for details) were referred to as oscillatory. When we considered the whole bandwidth from 1 to 100 Hz the fraction of oscillatory neurons did not differ significantly in TS and PD (36.15% vs. 26%, Fisher’s exact test, P = 0.30) with a majority of non-oscillatory neurons in both groups (Fig. 2A).

**Figure 2.**
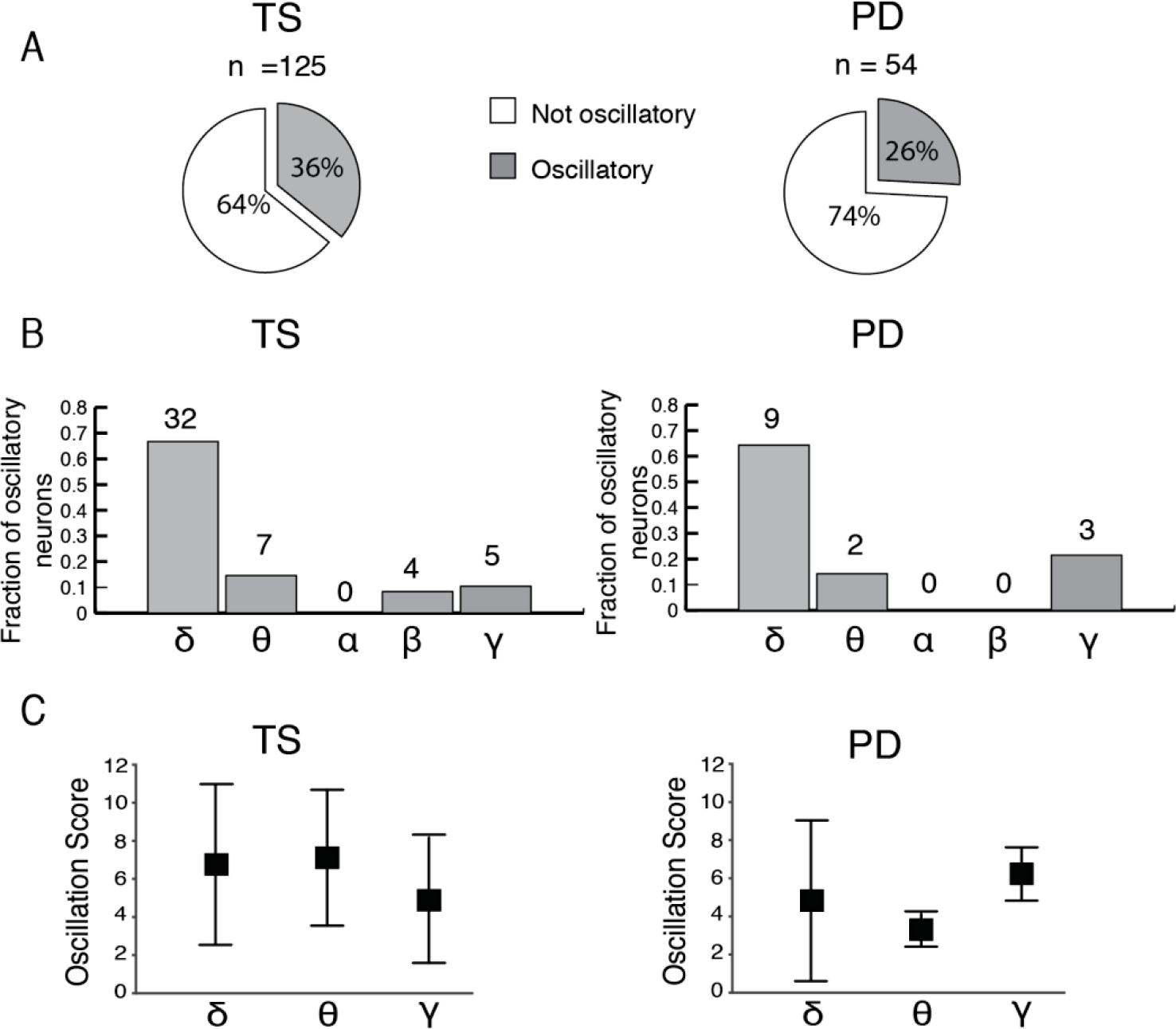
Single neuron oscillations in STN of TS and PD patients. Spectral analysis of STN neurons activity in TS (left column) and PD (right column) patients. *A*, Proportion of oscillatory (gray) and not-oscillatory (white) neurons in TS (N=125 neurons, left) and PD (N=54 neurons, right) groups. *B*, Distribution of oscillatory neurons across different frequency bands: *δ* (1-4) Hz, *θ* (4-8) Hz, *α* (8-12) Hz, *β* (12-30) Hz and *γ* (30-100) Hz in patients with TS (left) and PD (right). *C*, Oscillation strength for the relevant frequency bands (*δ, θ* and *γ*) for TS (left) and PD (right) groups.

Next, we analyzed whether the specific oscillatory content was similar. Table 3 summarizes the distribution of peak frequency observed in oscillatory neurons across the delta (1-4 Hz), theta (4-8Hz), alpha (8-12Hz), beta (12-30Hz) and gamma (30-100Hz) bands. Oscillations in the low-frequency and in the high-frequency bands modulation were prominent in both groups (δ: TS 64.3% vs. PD 64.3%; θ: TS 14.2% vs. PD 14.6%; γ: TS 21.5% vs. PD 10.4%) while alpha-band oscillations were absent in both TS and PD groups (Fig. 2B). The only barely significant difference was the presence of beta oscillations in TS and not in PD (n = 4; 8.3% vs n=0, unpaired test between percentages, P = 0.0485). However, the overall distribution of peaks across frequency bands did not differ between TS and PD (Fisher’s exact test, P = 0.59). We also tested for discrepancies in the oscillation strength between the groups comparing the oscillation scores in alpha, theta and gamma bands (Fig. 2C). Again, we did not find any specificity in the TS group (TS vs. PD; δ: 6.88±2.1 vs. 4.7±2.1, Mann-Whitney, P = 0.13; θ: 7.08±1.7 vs. 3.46±0.1, Mann-Whitney, P = 0.11; γ: 5.11±0.8 vs. 6.20±1.7, Mann-Whitney, P = 0.25). Our results show that TS STN activity does not show any specificity in the overall intensity of the firing rate or its dominant frequency bands. However, this might be due to a difficulty of these two coarse measurements to capture the marks of TS.

**Table 3.**
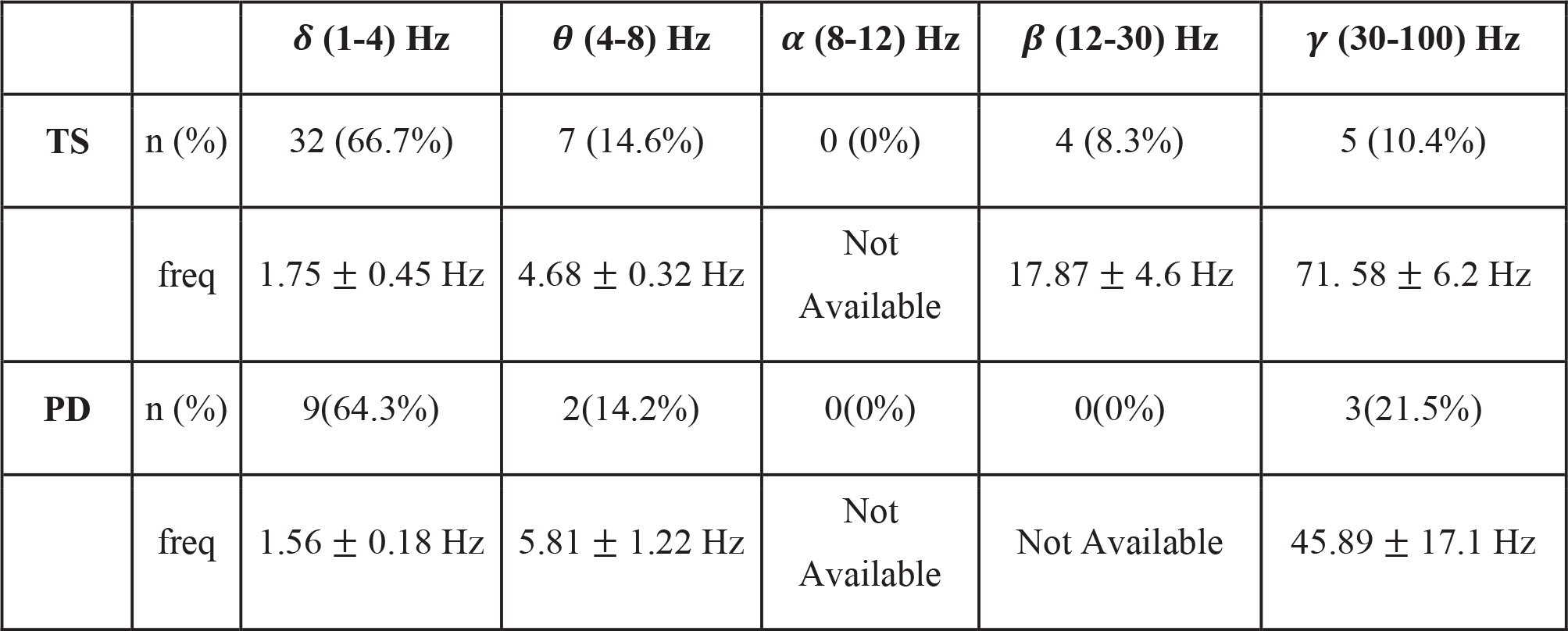
Oscillation frequencies of STN cells (median ± iqr/2) in TS and PD groups

### Slow bursting activity characterizes STN neurons in TS

We focused then on the temporal structure of the bursting activity which is known to be strongly related to TS tics. In particular, as neural activity underlying tics appear prevalently as arrhythmic bursts (Bronfeld *et al.*, 2011;Bronfeld & Bar-Gad, 2013), their dynamics could not be directly reflected in the spectral density but could instead be visible only with a detailed analysis of the temporal patterns of firing. We tested whether bursting regime was prevalent in TS by means of the analysis of the ISI distribution. We found that these distributions can be fit by gamma functions, and that *log*(*κ*)(the shape factor) of the function is a highly efficient indicator of the level of regularity and in particular the presence of bursts (see Methods for details). According to this measurement, we classified each neuron within three temporal patterns of firing: burst-like *log*(*κ*) < 0, regular tonic *log*(*κ*) > 0.3, characterized by a very stable ISI, and an intermediate state of irregular firing (Fig. 3A-C).

**Figure 3.**
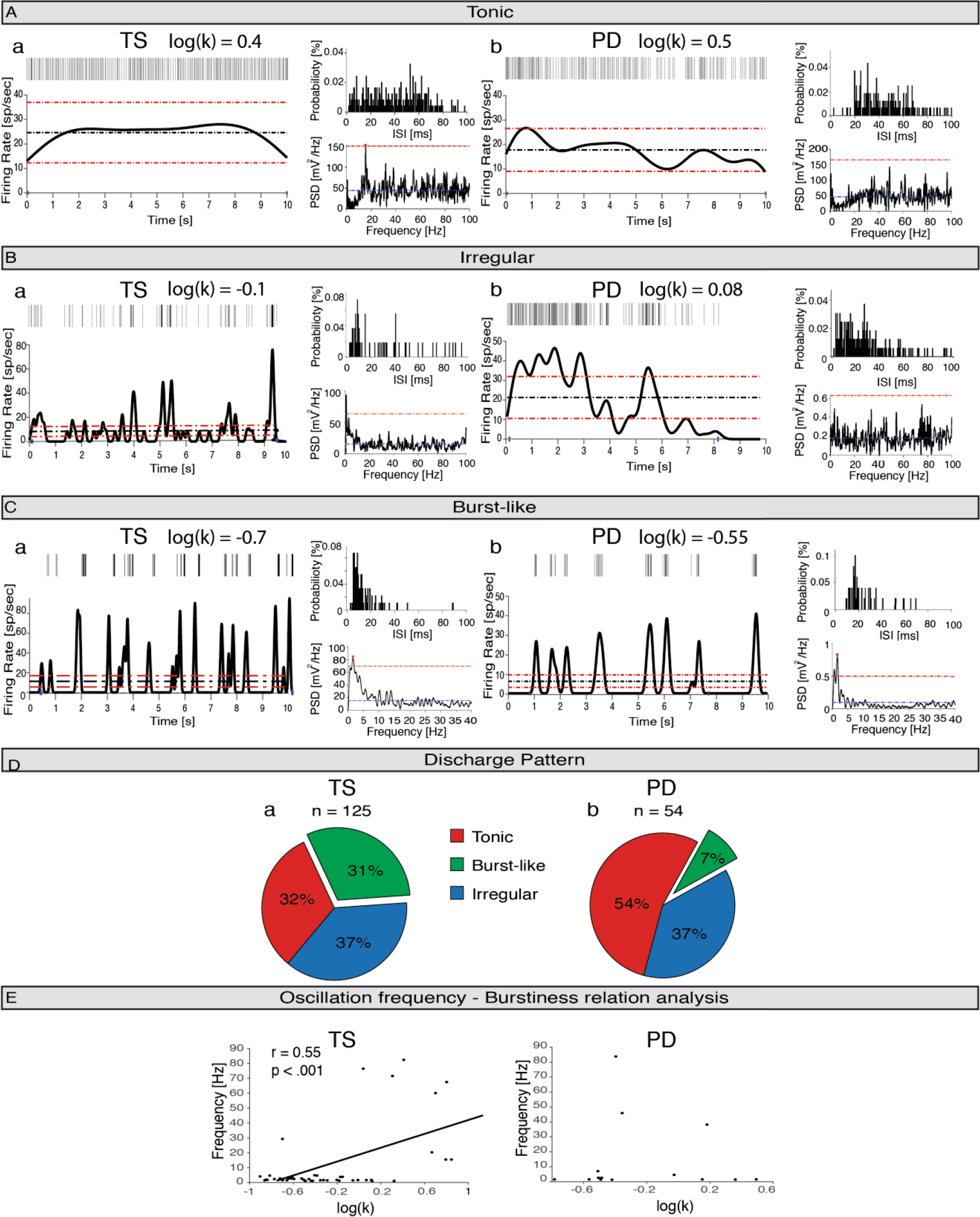
Firing patterns of STN neurons in TS and PD groups. *A-C* Representative temporal firing patterns of single units of STN in TS (Left, a column) and PD (Right, b column) patients: tonic firing (A), irregular firing (B) and burst-like firing (C) (see Methods for details). Each panel (Aa, Ba, Ca, Ab, Bb, Cb) is organized as follows: in the top left vertical bars indicate time stamps of trains of identified spikes; in the bottom left the firing rate of the neuron (tick black line), the mean firing rate (black dashed horizontal line) and the 0.5*mean range (red dashed horizontal lines); in the top right ISI probability distribution with 1 ms bin; in the bottom right the power spectrum density (black tick line), the mean power spectral density (blue dashed horizontal line) and the minimum threshold (red dashed horizontal line). *D*, Proportion of bursting (green), tonic (red) and irregular (blue) neurons in TS [Da] and PD [Db] groups. The number of the neurons analyzed in each group is displayed above the pie chart. *E*, Frequency of oscillation of the rhythmic neurons versus *log*(*κ*) in TS (left) and PD (right). A significant positive linear correlation was found in TS. Black thick line indicates the linear trend, while r and p are the Pearson correlation coefficient and the statistical significance respectively.

The distribution of the temporal structure of spiking activity in TS patients showed a large fraction of bursting neurons (39/125, 31.2%, Fig. 3D). In sharp contrast, in PD this fraction was only 7.4% (4/54) showing a significantly minor presence of bursting unit (F test, P <.001).

To assess whether TS and PD STN neurons differ in their relative temporal pattern distribution independently of the pattern categorization, the *log*(*κ*) was compared across all neurons of the two groups. The shape factor in TS STN neurons was negative (*log*(*κ*) = −0.3 ± 0.3) indicating a prevalence of bursting activity. The distribution of the shape factor was instead centered close to zero in PD STN neurons (*log*(*κ*) = −0.06 ± 0.33) with a significant difference relative to TS neurons (Mann-Whitney, P < 0.05). These results demonstrate that TS STN neurons operate closer to a burst-like regime in consistence of what inferred from spiking pattern classification.

Moreover, as shown in Table 4, the intra-burst firing rate of neurons with burst-like discharge in TS was really high TS: 176.3 ± 18 spikes/sec and significantly higher than the firing rate of burst-like neurons in PD (PD:107 ± 24 spikes/sec, Mann-Whitney test, P = 0.0406). No significant changes were noted instead in the other burst characteristics such as the average inter burst interval (IBI: 597±264 ms vs. 593 ± 207 ms, Mann-Whitney, P = 0.62) and its coefficient of variation (IBI CV: 1.15 ± 0.26 vs. 0.89 ± 0.07, Mann-Whitney, P = 0.14). Also, the number of spikes within bursts (SWB) and the fraction of intra bursts (SIB) spikes were found not to differ significantly (SWB: 4.17 ±0.35 vs. 3.77 ± 0.19, Mann-Whitney, P = 0.11, SIB: 52.43 ± 6.5 % vs. 53.34 ± 3.5 %, Mann-Whitney, P = 0.97). Overall, these results show that TS neurons are characterized by a large fraction of intensely bursting neurons.

**Table 4.** Bursts characteristics of bursting cells (median ± iqr/2) in TS and PD groups: Intra-burst firing rate (IBF), Inter-burst interval (IBI), number of spikes within bursts (SWB), percentage of spikes in bursts (SIB) and the coefficient of variation of the inter-burst interval (IBI CV).

|  | **TS** | **PD** | **P-value** |
| --- | --- | --- | --- |
| **IBF** | 176.3 ± 18 Hz | 107 ± 24 Hz | 0.0406* |
| **IBI** | 597± 264 ms | 593 ± 207 ms | 0.62 |
| **SWB** | 4.17 ± 0.35 | 3.77 ± 0.19 | 0.11 |
| **SIB** | 52.43 ± 6.5 % | 53.34 ± 3.5 % | 0.97 |
| **IBI CV** | 1.15 ± 0.26 | 0.89 ± 0.07 | 0.14 |

We wondered then whether these firing patterns were related to the presence of oscillatory neurons (see previous subsection). When stratifying by temporal pattern of firing, we found that in oscillatory neurons ranging from delta to alpha band oscillation bursting characteristics were prevalent (n = 31/50; 62%), whereas beta and gamma oscillations were exclusively found in not-burst-like neurons (n = 12/12; 100%). Moreover, we found among TS neurons a significant positive correlation between the shape factor *log*(*κ*) and the frequency of oscillation in the oscillatory cells in (Pearson’s coefficient r = 0.55, P < .001) (Fig. 3E). This implies that, although low frequency oscillations are present in both PD and TS STNs, only in TS they are specifically originated by bursting activity.

### TS bursting neurons are prevalently located close to clinical DBS target

Microelectrode recordings were performed to locate the optimal depth for DBS stimulation. We investigated then if the neuron activity classification described above varied across depths and whether it was related to the location of optimal DBS target depth.

Neurons were recorded with 0.5 mm intervals from −6 to 4 mm from the target reference frame (see Methods for details). No significant linear correlation between firing rate and depth was observed (TS: Pearson’s coefficient r = −0.2, P = 0.07, PD: Pearson’s coefficient r = .32, P = 0.06). Distribution of FR across depths was flat in both groups and the lack of significant difference between FR of PD and TS neurons held true layer-wise (P> 0.23 for all depths Fig. 4A).

**Figure 4.**
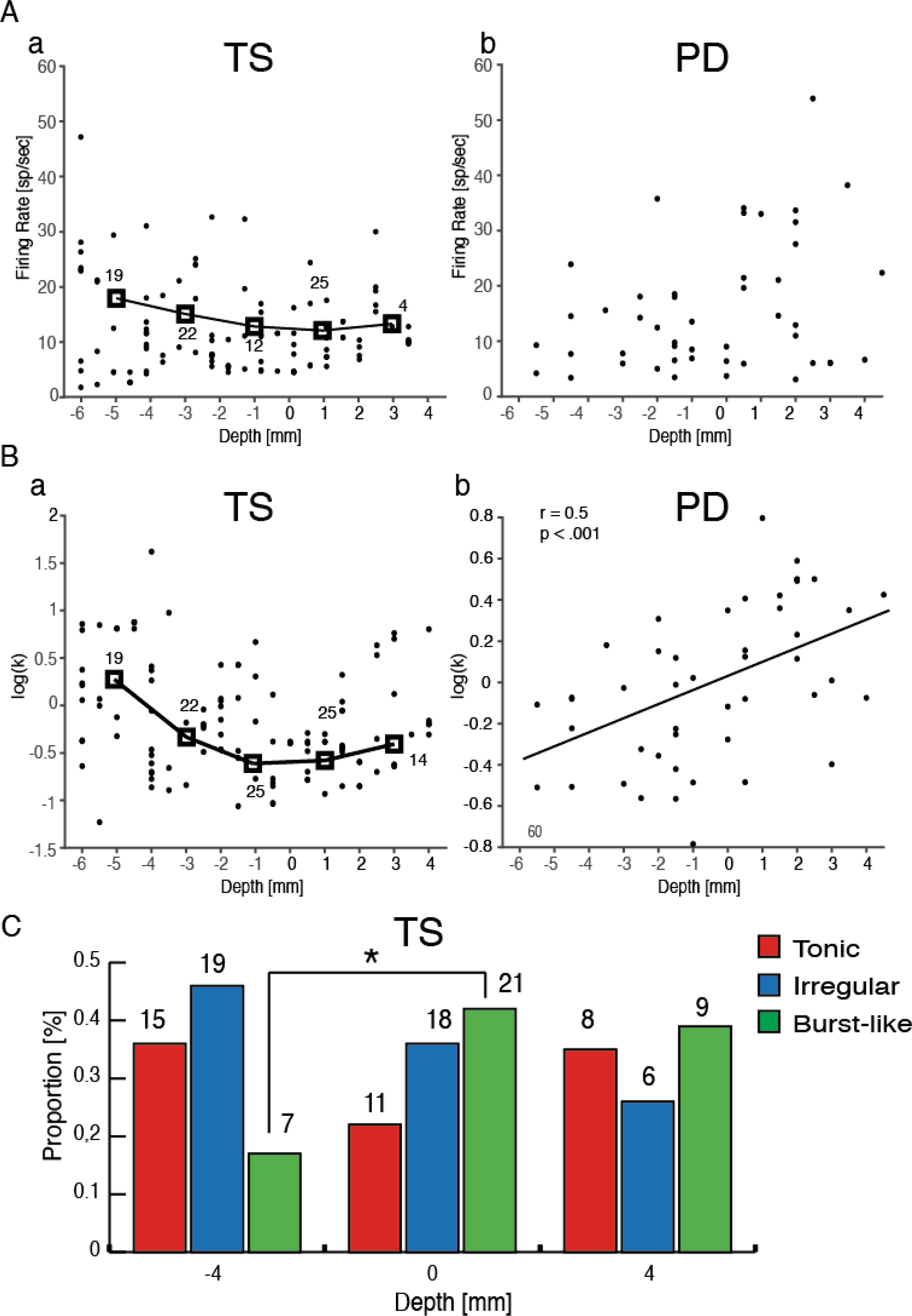
Topographical analysis of discharge patterns of STN neurons in TS and PD. TS patients results are displayed on the left, (a column) and PD patients results on the right (b column). *A*, distribution of firing rate as a function of the distance, ranging from −6 mm (dorsal) to 4 mm (ventral). The 0 mm depth is defined as the depth of the selected target for DBS. Black thick squares indicate the mean of 2 mm segments. The number of neurons included in each group is reported next tothe corresponding bin. *B*, same as A for *log*(*κ*). r and p are the Pearson correlation coefficient and the associated significance respectively. Black thick line indicates the linear trend. *C*, Distribution of TS neurons according to spiking pattern (Tonic (red), Irregular (blue) and bursting neurons (green)) and target relative electrode depth (around −4 mm, 0 mm and 4 mm). Asterisks indicate significant difference between bursting neurons proportions at −4 mm and 0 mm depth (Fisher Exact test, P = 0.01).

Oscillatory activity over all frequencies was also equally distributed throughout the STN, but it was found that gamma oscillatory neurons were exclusively found before and after the target portion along the dorsal-ventral trajectory in TS (relative target-depth: < −2 mm and > 2 mm).

The topographical analysis for *log*(*κ*) excluded a linear relationship between burstiness and depth in TS group (Pearson’s coefficient r = −0.16, P = 0.09). The shape factor varied instead as a function of depth in a U-shaped way (Fig. 4B): bursting neurons were found more frequently close to the selected DBS target (mean depth: −0.93 mm, SD: 2 mm). This is radically different from results in PD patients in which the shape factor grew linearly with the depth (Pearson’s coefficient r = 0.5, P < 0.001) and the mean depth of bursting neurons was −2 mm (mean −2 mm; SD: 3.5 mm) Moreover, we found a non-homogeneous spatial distribution across depth of the pattern discharge of TS neurons (Fisher Exact test, P = 0.03) with a maximum bursting neurons prevalence at target depth (42% of bursting neurons at depth = 0mm, vs 17% bursting neurons at depth −4mm, post-hoc Fisher Exact test, P = 0.01) (Fig 4C).

This suggests that in TS STN presents a global bursting activity over all depths, but with a significantly higher density close to the optimal clinical DBS target.

Finally, we computed the amount of information about the target depth carried by the presence of bursting neurons, opposed to presence of oscillatory or high firing neuron. We found that the firing rate did not carry any information, while both the presence of bursting neurons or oscillatory neurons carried significant information (P<0.05, bootstrap test, see Methods for details), with the presence of bursts carrying 12% more information than the presence of oscillations. This corroborates the idea that the recording of bursting neurons can help in the location of the optimal DBS target.

## Discussion

Four TS patients were implanted with DBS in the STN and showed a significant improvement in motor disorders. We investigated the spatial and temporal structure of STN activity recorded from intraoperative DBS surgery microrecordings under general anesthesia, analyzing depth-wise the firing activity of single neurons. STN activity of anesthetized PD patients was analyzed only for comparison. We found several results indicating that STN in TS patients is characterized by an arrhythmic temporal structure of neuronal activity. Indeed, we observed that firing rate in the TS STN was not significantly different from that in PD, suggesting that a rate description of basal ganglia is not sufficient to capture the differences between the two disorders. Also, from the point of view of spectral analysis we found no TS-specific hallmark, neither in the fraction of oscillatory neurons, nor in the preferred bands, nor in the power associated to each band (Fig. 2). TS-specific features became apparent only when we analyzed specifically bursting activity (Fig. 3). First, STN neurons in TS patients were to a larger extent exhibiting bursting activity, and a stronger activity within bursts. Second, bursting activity displayed a characteristic temporal scale (in the delta range) and a depth preference coherent with optimal location for clinical practice.

This is to our knowledge the first study on STN subterritories on TS in human patients. The main result is that STN in TS is characterized by low frequency irregular bursting neurons, and that these neurons cluster close to the anatomical location of the contacts whose stimulation produced the wider therapeutic window to control tics.

### Functional architecture of STN and motor disorders

Dynamics of firing of the basal ganglia in TS has been analyzed so far, but mainly in animal models, and usually with a specific lack of analysis of STN firing (Bronfeld *et al.*, 2011;Bronfeld & Bar-Gad, 2013;Yael *et al.*, 2015). Consequently, our work is the first to our knowledge able to test theoretical predictions of a recent broad computational model of Tourette syndrome (Caligiore *et al.*, 2017). We note that model predicts burst-like activation of STN with an intra-burst frequency (IBF) of about 200 Hz that is close to the level of firing we observed in our recordings.

Recent studies performed similar analysis on subterritories of the STN in PD patients. A clinical paper (Pozzi *et al.*, 2016) in particular found spatial differences in bursting activities across functional domains in the STN in awake PD patients. Here we focused on an anatomical analysis mainly within a single functional domain (sensory-motor, see Fig. 1) and we found depth specific bursting properties. Although the two works investigate distinct conditions with different statistical analysis, together they strongly suggest that STN dynamics might be far less homogeneous than generally thought. This has obvious implications in the relevance of an accurate localization of DBS stimulation (Montgomery, Jr., 2012). Moreover, such localization of bursts might be, through the hyper-direct pathway, among the causes of the localization of bursts observed in the thalamus (Marceglia *et al.*, 2010), which in turn is supposed to be associated to the specific location of the tics.

### STN DBS for TS therapy

The main limitation of the current study is the dataset composed by only four patients. However even such a small set is significant when compared with the lack of similar data in literature. A recent meta-analysis on DBS in TS patients (Baldermann *et al.*, 2016) reported no patients with STN implant. To the best of our knowledge, the only previous report of efficacy of STN DBS for tics was observed in a PD patient with concomitant tourettism (Martinez-Torres *et al.*, 2009). Our dataset is then the first relevant insight into the dynamics of STN in TS patients. Moreover, thanks to the quality of our recordings we were able to record from a large number of neurons (n=125).

### STN dynamics and anesthesia

To which extent the behavior we observe in TS STN is due to anesthesia? Even though almost all PD patients are implanted without anesthesia, we found two anesthetized PD patients to control for the effects of anesthesia. Although this dataset is limited, we were in the end able to collect a pool of 54 neurons, and the results we report constitute the best available control for the behavior of the anesthetized STN in TS patients. Moreover, with the current state of DBS implant technology, implants in TS patients will be always performed under anesthesia to avoid damages due to the tics of the patients and the STN activity will be recorded in the same conditions analyzed in this work; hence the presence of bursts might be able to help in the localization of the preferred depth for clinical purposes in any practical clinical situation. The main limitation of anesthesia is instead that it prevents us from answering is whether the observed low frequency intense bursts are correlated to the onset of tics, as observed in animal models (Bronfeld *et al.*, 2013).

### Role of STN in basal ganglia dysfunctions underlying TS

As mentioned in the introduction, TS disorders are thought to originate from a striatum-driven overactivation of the DP and hence to a strong excessive positive feedback to the cortex leading to involuntary movements (Bronfeld *et al.*, 2011;Bronfeld & Bar-Gad, 2013;Ganos *et al.*, 2015;Mink, 2001b;Mink, 2006;Yael *et al.*, 2015). As STN does not take part to DP, this model does not account for the STN dynamics observed in this work. The observed bursts, however, might be originated by the STN excitation following striatum overactivation through the IP. In this case, a scattered and non-effective IP inhibition might contribute to tics by preventing an accurate termination of the movements. An STN on burst mode, however, should affect also the HP, which in the selection model is thought to reset the activity of the thalamus to prepare for the activation through DP of the desired movement. In this case, an abnormal temporal distribution of the STN activity might again result in an insufficient inhibition of the thalamus, laying the ground for tics onset. Overall, our results hint toward the necessity of developing a broader model of TS origin, including all three loops, and provide some first pieces of evidence to achieve this result.

## Acknowledgment

Authors wish to acknowledge Dr. Francesco Carella, for his aid in microrecording experimental design and results interpretation. S.M. was supported by the Bertarelli Foundation and by institutional funds from Scuola Superiore Sant’Anna and École Polytechnique Fédérale de Lausanne. L.M.R. was partly supported by the Italian Ministry of Health grant GR-2009-1594645. A.M. was supported by institutional funds from Scuola Superiore Sant’Anna.

